# *osr1* couples intermediate mesoderm cell fate with temporal dynamics of vessel progenitor cell differentiation

**DOI:** 10.1101/2020.11.14.383141

**Authors:** Elliot A. Perens, Jessyka T. Diaz, Agathe Quesnel, Amjad Askary, J. Gage Crump, Deborah Yelon

## Abstract

Transcriptional regulatory networks refine gene expression boundaries throughout embryonic development to define the precise dimensions of organ progenitor territories. Kidney progenitors originate within the intermediate mesoderm (IM), but the pathways that establish the boundary between the IM and its neighboring vessel progenitors are poorly understood. Here, we delineate new roles for the zinc finger transcription factor Osr1 in kidney and vessel progenitor development. Zebrafish *osr1* mutants display decreased IM formation and premature emergence of neighboring lateral vessel progenitors (LVPs). These phenotypes contrast with the increased IM and absent LVPs observed with loss of the bHLH transcription factor Hand2, and loss of *hand2* partially suppresses the *osr1* mutant phenotypes. *hand2* and *osr1* are both expressed in the posterior lateral mesoderm, but *osr1* expression decreases dramatically prior to LVP emergence. Overexpressing *osr1* inhibits LVP development while enhancing IM formation. Together, our data demonstrate that *osr1* modulates both the extent of IM formation and the temporal dynamics of LVP development, suggesting that a balance between levels of *osr1* and *hand2* expression is essential to demarcate the dimensions of kidney and vessel progenitor territories.

**SUMMARY STATEMENT:** Analysis of the *osr1* mutant phenotype reveals roles in determining the extent of intermediate mesoderm formation while inhibiting premature differentiation of neighboring vessel progenitors.

## INTRODUCTION

Proper embryonic patterning depends on the establishment of progenitor territories with well-defined gene expression patterns (Briscoe and Small, 2015). For example, the medial-lateral axis of the vertebrate posterior mesoderm is divided into precise stripes of progenitor territories that give rise to various organs and cell types, including the kidneys, blood vessels, and blood cells (Prummel et al., 2020). Kidney progenitors originate within a pair of bilateral territories called the intermediate mesoderm (IM) (Davidson et al., 2019; Dressler, 2009; Gerlach and Wingert, 2013). While transcription factors such as Lhx1/Lim1 and Pax2 are known to be expressed in the IM and required for its development (Barnes et al., 1994; Carroll and Vize, 1999; Carroll et al., 1999; Cirio et al., 2011; Dressler et al., 1990; Torres et al., 1995; Tsang et al., 2000), the mechanisms that establish boundaries between the IM and its neighboring territories remain poorly understood.

The zinc finger transcription factor Osr1 is an intriguing candidate for playing a central role in IM boundary formation. Gene expression analyses and temporal fate mapping in amniotes demonstrated that *Osr1* is expressed initially in the IM and the neighboring lateral plate mesoderm, which contains vessel progenitors, before its expression becomes restricted to kidney progenitors (James et al., 2006; Mugford et al., 2008). In zebrafish, *osr1* is expressed in the lateral posterior mesoderm, initially adjacent to the IM; later, a stripe of lateral vessel progenitors (LVPs) arises between the IM and the *osr1*-expressing territory (Mudumana et al., 2008; Perens et al., 2016). Mouse *Osr1* knockout embryos have decreased *Lhx1* and *Pax2* expression during early stages of kidney development, thought to be due to increased apoptosis (Wang et al., 2005). In zebrafish, *osr1* knockdown studies yielded varying conclusions: one study of *osr1* morphants determined that *osr1* is only required for maintenance of the pronephron lineage (Mudumana et al., 2008), while another indicated that *osr1* was required for IM formation (Tena et al., 2007). Additionally, *osr1* knockdown resulted in an expanded venous vasculature (Mudumana et al., 2008). Because LVPs are involved in vein formation (Kohli et al., 2013), it is interesting to consider whether *osr1* may influence LVP development. In total, however, the functions of *osr1* in the initial development of the IM and vessel progenitors remain unclear.

Considering the discrepancies often seen between morphants and mutants (Schulte-Merker and Stainier, 2014), we chose to augment prior *osr1* morphant studies (Mudumana et al., 2008; Tena et al., 2007; Tomar et al., 2014) by analyzing a TALEN-generated *osr1* mutation (Askary et al., 2017). The *osr1* mutant phenotype demonstrated important roles of *osr1* in promoting IM and pronephron differentiation and inhibiting premature LVP formation. Previously, we found that *osr1* and *hand2*, which encodes a bHLH transcription factor, are co-expressed in the lateral posterior mesoderm and that *hand2* promotes LVP development while inhibiting the lateral extent of IM formation (Perens et al., 2016). Each of these phenotypes was partially suppressed by mutation of *osr1*. Intriguingly, wild-type embryos displayed a striking reduction of *osr1* expression just prior to LVP formation, while overexpression of *osr1* inhibited LVP emergence while elevating IM formation. Together, our studies suggest a new model in which *osr1* expression dynamics balance IM differentiation with the temporal emergence of neighboring vessel progenitors.

## RESULTS AND DISCUSSION

### Mutation of *osr1* has graded effects along the proximal-distal axis of the pronephron

To enhance our understanding of *osr1* function in the posterior mesoderm, we analyzed a TALEN-generated *osr1* mutation (Askary et al., 2017). *osr1^el593^* is a seven base pair deletion leading to a frameshift; the predicted mutant Osr1 protein would contain its first 80 amino acids, followed by 45 missense amino acids, and would lack its zinc fingers (Fig. 1A). Thus, *osr1^el593^* is likely a strong loss-of-function allele.

**Figure 1.**
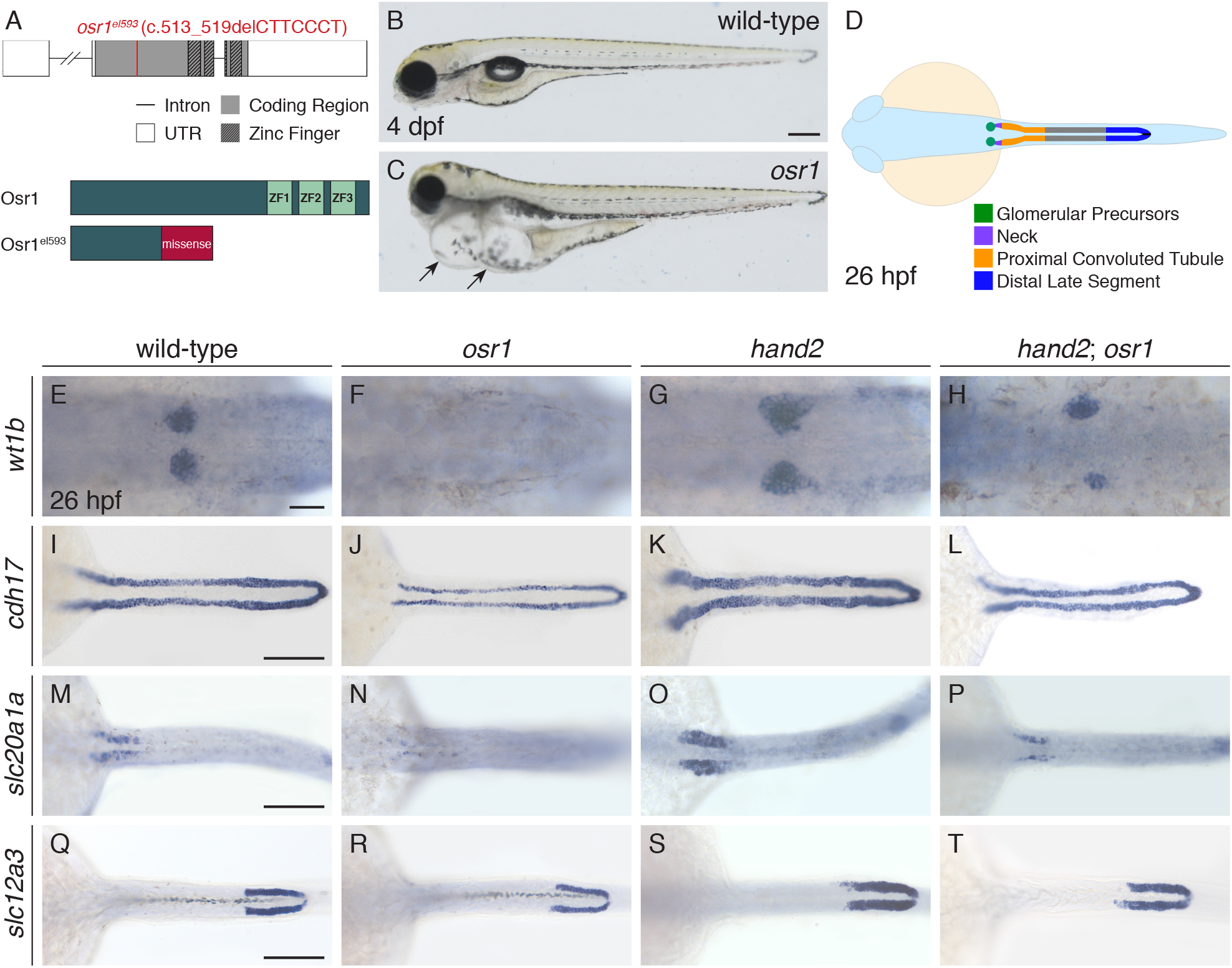
Pronephron defects in *osr1* mutants are partially suppressed by *hand2* loss-of-function. (A) *osr1^el593^* is a TALEN-generated 7 bp deletion allele. Schematics show gene structure with location of deletion and predicted wild-type and Osr1^el593^ proteins (ZF, zinc finger). (B,C) Lateral views, anterior to the left, of wild-type (B) and *osr1* mutant (C) embryos at 4 days post-fertilization (dpf). *osr1* mutants display severe pericardial and body wall edema (arrows). (D-T) Dorsal views, anterior to the left, of a pronephron schematic (D), wild-type (E,I,M,Q), *osr1* mutant (F,J,N,R), *hand2* mutant (G,K,O,S), and *hand2*; *osr1* double mutant (H,L,P,T) embryos at 26 hours post-fertilization (hpf). In situ hybridization shows expression of *wt1b* (E-H) in glomerular precursors, *cdh17* (I-L) throughout the tubules, *slc20a1a* (M-P) in the proximal convoluted tubules, and *slc12a3* (Q-T) in the distal late segments. Compared to wild-type (E,I,M,Q), expression is absent (F), thin and shortened anteriorly (J), reduced (N), and thin (R) in *osr1* mutants; expanded in *hand2* mutants (G,K,O,S); and relatively similar to wild-type in *hand2*; *osr1* double mutants (H,L,P,T). Scale bars: 200 μm (B,C), 100 μm (I-T), 25 μm (E-H).

Homozygous *osr1* mutant embryos exhibited progressive pericardial and body edema (Fig. 1B,C) comparable to other zebrafish mutants with defects in pronephron development (Kroeger et al., 2017; Lun and Brand, 1998). Consistent with this observation, *osr1* mutants displayed deficits in multiple pronephron segments. Most dramatically, markers of the glomerular precursors (*wt1b*, Fig. 1F) and glomerular precursors and neck region (*pax2a*, Fig. S1B), were absent. *cdh17*, which is normally expressed throughout the pronephron tubules (Fig. 1I), lacked expression at the proximal extent of the tubule, while the remaining tubule appeared thinner throughout (Fig. 1J). Similarly, a marker of the proximal convoluted tubule segment (*slc20a1a*, Fig. 1M,N) was reduced in intensity, while a marker of the distal late segment (*slc12a3*, Fig. 1Q,R) revealed a slightly thinner expression pattern in the mutant. Thus, while prior morphant studies found that *osr1* is only required for proximal tubule development (Mudumana et al., 2008), the *osr1* mutant phenotype revealed that *osr1* is required for proper development of the entire pronephron but has a higher impact on the development of the proximal territories.

The strong proximal deficiencies observed in *osr1* mutants were reminiscent of the pronephron phenotypes previously shown to result from overexpression of *hand2* (Perens et al., 2016). Additionally, pronephron size was increased in a *hand2* null mutant (Perens et al., 2016) (Fig. 1G,K,O,S), and knockdown of *osr1* function has been shown to partially suppress this *hand2* mutant phenotype (Perens et al., 2016). Confirming this genetic interaction, we found that mutation of *osr1* also partially suppressed the pronephron phenotypes in *hand2* mutants (Fig. 1H,L,P,T; Fig. S1D).

### *osr1* is required to generate the full complement of intermediate mesoderm

Previously we found that *hand2* constrains the size of the pronephron by repressing IM formation (Perens et al., 2016). Considering the genetic interaction between *hand2* and *osr1* during pronephron development, we investigated whether *osr1* also regulates the extent of initial IM formation. In *osr1* mutants, we observed narrowed expression of *lhx1a* in the IM compared to that seen in wild-type embryos (Fig. 2A,B). Quantification of the number of Pax2a^+^ cells revealed that this altered IM appearance reflected a significant decrease in the number of IM cells in *osr1* mutants (Fig. 2I). Thus, while *osr1* knockdown studies left unclear whether *osr1* has an early role in IM formation (Mudumana et al., 2008; Tena et al., 2007), the *osr1* mutant phenotype indicated that *osr1* is required for the initial generation of the full complement of IM cells. This phenotype also raises the possibility that the IM is divided into *osr1*-dependent and *osr1*-independent subregions. Furthermore, we found that loss of *hand2* function partially suppressed the IM defects in *osr1* mutants (Fig. 2D,H,I). Thus, *osr1* and *hand2* operate in antagonistic genetic pathways to regulate IM differentiation, raising the possibility that *osr1*, like *hand2*, may execute this function at the lateral border of the IM.

**Figure 2.**
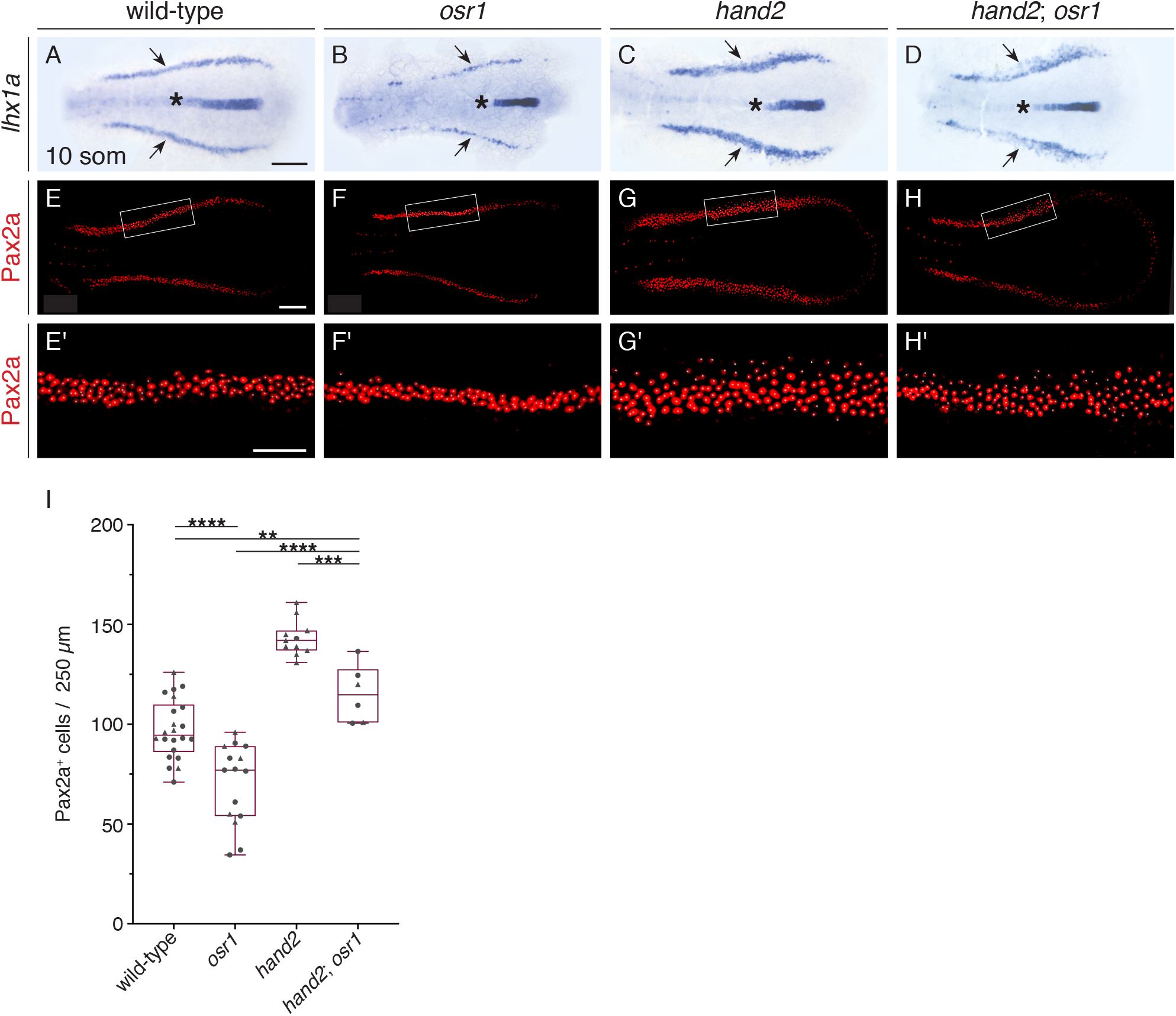
*osr1* is required to promote intermediate mesoderm differentiation. (A-H) Dorsal views, anterior to the left, of the posterior mesoderm at the 10 somite stage (som). (A-D) In situ hybridization shows expression of *lhx1a* in the IM (arrows). Compared to wild-type (A), expression is narrowed in *osr1* (B), widened in *hand2* (C), and irregular in *hand2*; *osr1* (D) embryos. Expression in the notochord (asterisk) is unaffected. Unlike the width of the IM, we did not observe a change in the proximal-distal length of the IM in mutant embryos. (E-H) Three-dimensional reconstructions of Pax2a immunofluorescence in the IM of wild-type (E), *osr1* (F), *hand2* (G), and *osr1*; *hand2* (H) embryos. (E’-H’) Magnification of boxed 250 μm long regions used for counting Pax2a^+^ cells. White dots indicate Pax2a^+^ nuclei. (I) Quantification of Pax2a^+^ cells in a 250 μm long portion of the IM in the indicated genotypes. Symbols represent individual embryos (circles, average of left and right IMs; triangles, single IM; see Materials and Methods); boxes represent interquartile range; central line marks the median; whiskers indicate maximum and minimum values. *P* values calculated using non-parametric Mann-Whitney U tests: **** *P*<0.0001, *** *P*=0.0006, ** *P*=0.0094. Scale bars: 100 μm (A-H), 50 μm (E’-H’).

### *osr1* inhibits the premature emergence of lateral vessel progenitors in the posterior mesoderm

We investigated whether *osr1*, like *hand2*, might play a role in the development of the neighboring lateral vessel progenitors (LVPs). The LVPs normally arise at the lateral boundary between the IM and the *hand2*/*osr1*-expressing territory of the posterior mesoderm during a short, consistent time window (Kohli et al., 2013; Perens et al., 2016) (Fig. 3A,B,E). Surprisingly, however, we found that LVPs form prematurely in *osr1* mutants. Specifically, while *etv2*-expressing LVPs rarely appeared prior to 10 som in wild-type embryos, some *osr1* mutants exhibited *etv2*-expressing LVPs as early as 6 som, and most *osr1* mutants possessed *etv2*-expressing LVPs by 8 som (Fig. 3C-E, Fig. S2). Additionally, *etv2* expression in a separate proximal region, which has a morphological arrangement distinct from the more posterior territories (Fig. 3B,F), was increased in *osr1* mutants (Fig. 3B,D,F,G; Fig. S2). Thus, *osr1* is required to constrain vessel progenitor development, including inhibition of the premature differentiation of LVPs at the lateral border of the IM.

**Figure 3.**
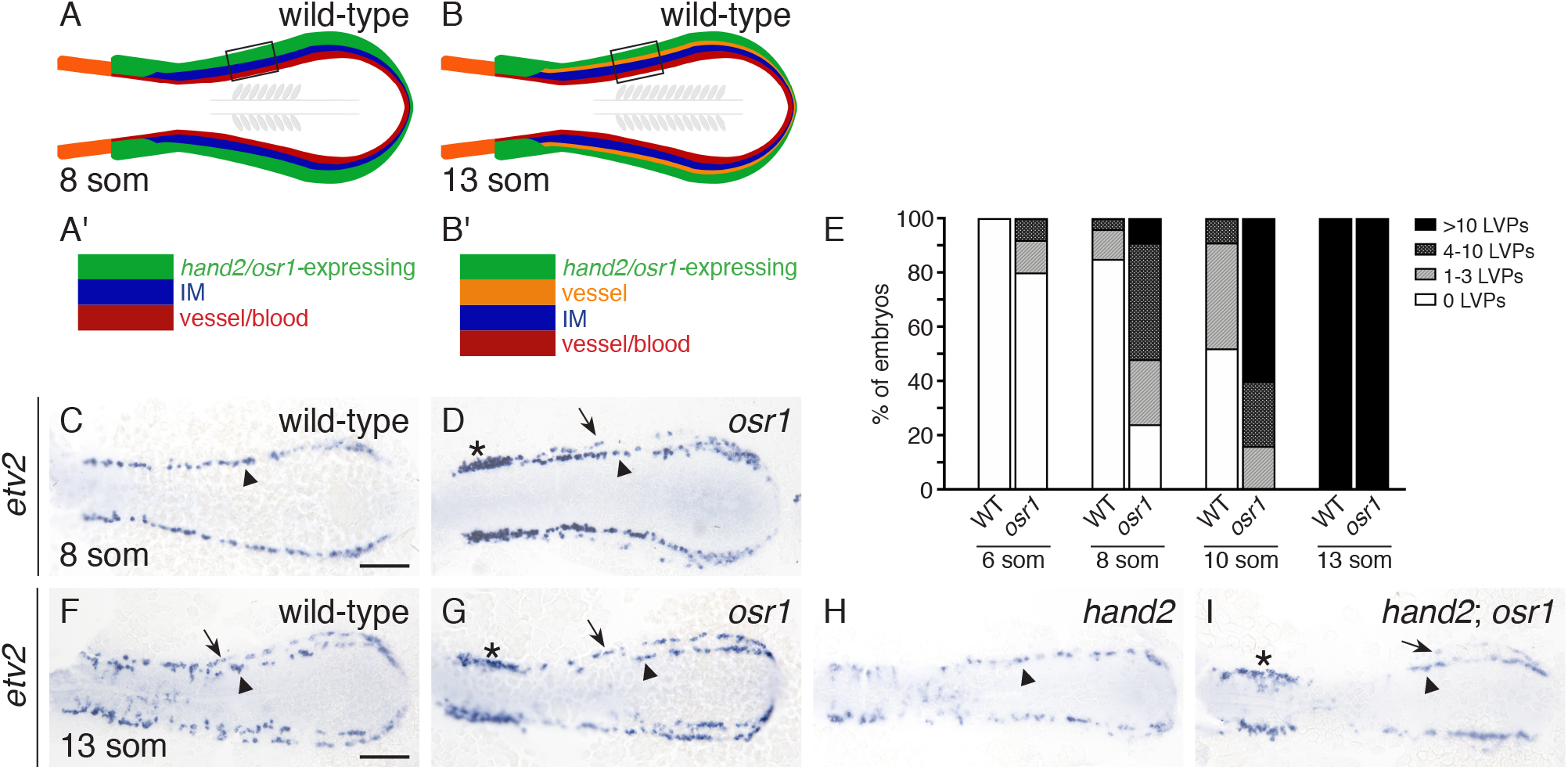
*osr1* inhibits the premature emergence of lateral vessel progenitors. (A,B) Schematics depict lateral posterior mesoderm territories, dorsal views, anterior to the left. (A’,B’) Expansion of corresponding boxed regions. (C,D,F-I) In situ hybridization shows *etv2* expression in wild-type (C,F), *osr1* (D,G), *hand2* (H) and *hand2*; *osr1* (I) embryos; dorsal views, anterior to the left, at 8 (C,D) and 13 (F-I) som. (C,D) At 8 som, *etv2* is expressed in a relatively medial territory (arrowhead) on each side of the wild-type mesoderm (C). In *osr1* mutants, *etv2* is also expressed in some relatively lateral cells (D, arrow), and its expression is increased proximally (asterisk). (E) Quantification of LVPs in wild-type and *osr1* mutant embryos at 6, 8, 10 and 13 som; sample sizes provided in Fig. S2E. Embryos were categorized based on the number of cells observed on whichever side of the mesoderm exhibited more LVPs. (F-I) At 13 som, *etv2* is expressed in both relatively medial (arrowheads) and lateral territories (arrows) on each side of wild-type (F), *osr1* (G) and *hand2*; *osr1* (I) embryos. In *hand2* mutants (H), *etv2* is expressed only in the medial territory. In *osr1* (G) and *hand2*; *osr1* (I) embryos, *etv2* expression is increased proximally (asterisk). Scale bars: 100 μm.

Because *osr1* and *hand2* interact antagonistically during IM development, we examined whether the same genetic interaction occurs during vessel progenitor development. Notably, while *hand2* mutants rarely form LVPs (Fig. 3H; 74% no LVPs, 26% 1-3 LVPs, n=31) (Perens et al., 2016), more *etv2*-expressing LVPs do form in *hand2*; *osr1* double mutants (Fig. 3I; 22% no LVPs, 56% 1-3 LVPs, 22% 4-10 LVPs, n=9). Taken together, these data suggest that, as in the IM, *osr1* and *hand2* act in antagonistic genetic pathways to regulate LVP formation.

In addition to the IM and LVP phenotypes, we found that *gata1*-expressing blood progenitors were reduced throughout the posterior mesoderm in *osr1* mutants (Fig. S3B). Unlike the IM and LVPs, however, blood progenitors did not seem altered in *hand2* mutants (Fig. S3C), and the *osr1* mutant phenotype did not appear to be suppressed by loss of *hand2* (Fig. S3D). Thus, in addition to the antagonistic pathways through which *osr1* and *hand2* regulate IM and LVP formation, *osr1* may function in additional genetic pathways that influence posterior mesoderm patterning. Other studies have implicated *osr1* in early mesendoderm development: *osr1* is expressed in mesendoderm prior to gastrulation, and *osr1* knockdown resulted in increased endoderm formation (Mudumana et al., 2008; Terashima et al., 2014). We, however, did not find an evident endoderm phenotype in *osr1* mutants (Fig. S4).

### *osr1* expression levels mediate intermediate mesoderm and lateral vessel progenitor cell fate decisions

Considering the dynamic nature of *Osr1* expression in amniotes (James et al., 2006; Mugford et al., 2008), we surmised that *osr1* expression in the zebrafish posterior mesoderm might be dynamic as well. Indeed, unlike *hand2*, which has consistent expression in the posterior mesoderm from tailbud stage to 10 som (Fig. 4A-C), *osr1* expression decreases dramatically during this same time period and prior to the emergence of the LVPs (Fig. 4D-F). Thus, decreased *osr1* expression may be required for LVP emergence.

**Figure 4.**
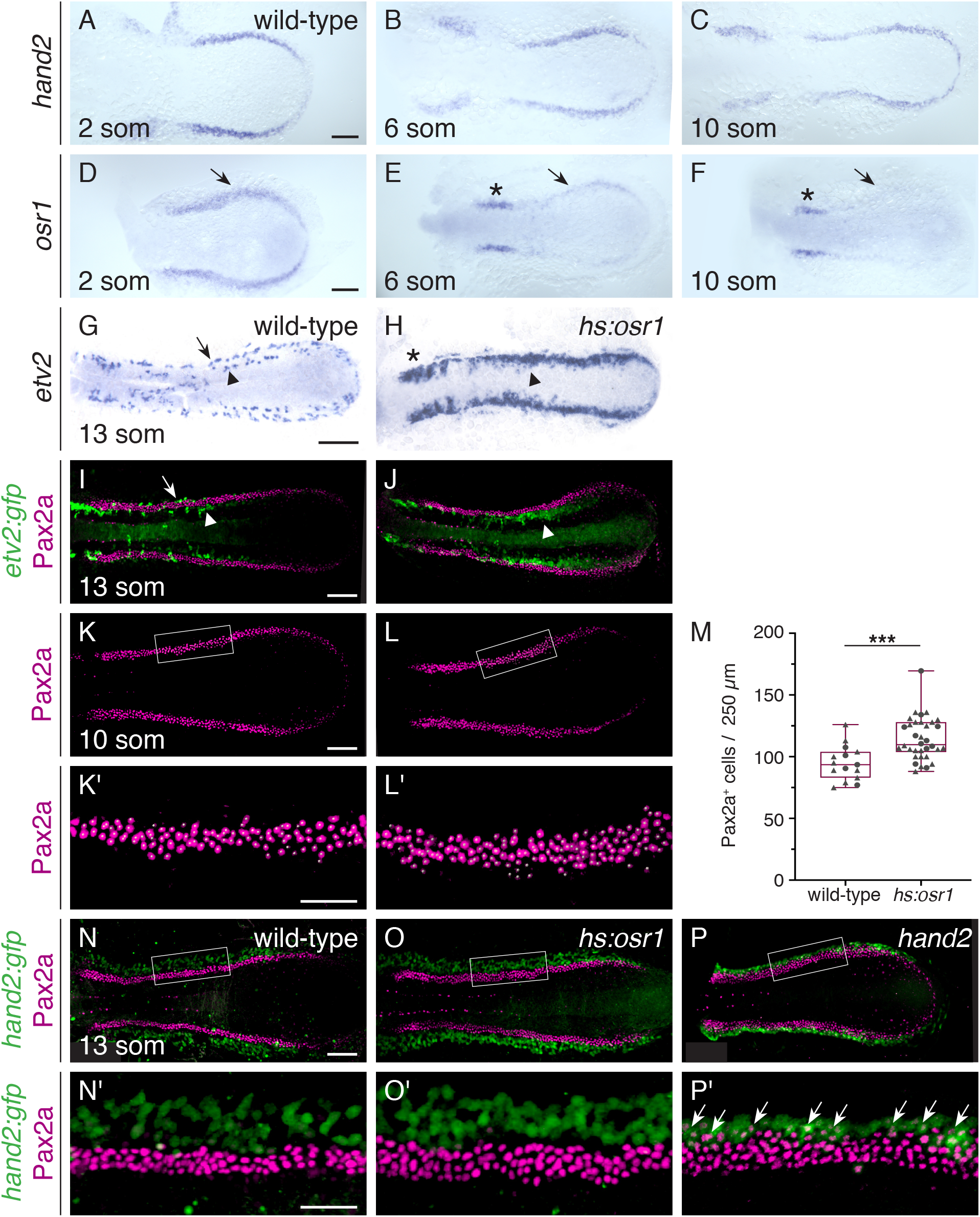
Increased *osr1* expression inhibits lateral vessel progenitor emergence while elevating IM formation. In situ hybridization (A-H) and immunofluorescence (I-L,N-P) indicate expression patterns in the posterior mesoderm; dorsal views, anterior to the left. (A-F) Between 2 and 10 som, *hand2* remains expressed in a relatively broad domain of the wild-type posterior mesoderm (A-C). In contrast, *osr1* expression is initially broad (D, arrow) but then becomes reduced in width and intensity (E,F; arrows), except proximally, where expression remains strong (asterisks), possibly representing expression in glomerular precursors (Tomar et al., 2014). (G,H) In contrast to wild-type embryos (G), which display *etv2* expression in both medial (arrowhead) and lateral (arrow) territories, *hs:osr1* embryos (H) display increased expression of *etv2* in medial (arrowhead) and proximal (asterisk) territories, but decreased *etv2* expression in lateral territories at 13 som. (I-L,N-P) Three-dimensional reconstructions of Pax2a and GFP immunofluorescence in wild-type (I,K,N), *hs:osr1* (J,L,O), and *hand2* mutant (P) embryos carrying *Tg(etv2:egfp)* (I,J) or *Tg(hand2:egfp)* (N-P). (K’,L’,N’-P’) Magnification of boxed 250 μm long regions; white dots indicate Pax2a^+^ nuclei (K’,L’). (I,J) While *etv2:egfp* expression is seen in both medial (arrowhead) and lateral (arrow) territories in wild-type at 13 som (I), *etv2:egfp* expression is only seen in a wide territory medial to the Pax2a^+^ IM in *hs:osr1* embryos (arrowhead). (K-M) Quantification of Pax2a^+^ cells in a 250 μm long portion of the IM, as in Fig. 2I, demonstrates a significant increase in IM cells in *hs:osr1* embryos. *P* value calculated using non-parametric Mann-Whitney U test: *** *P*=0.002. (N-P) Compared to wild-type (N) and *hs:osr1* (O) embryos, *hand2* mutant embryos (P) have a noticeable increase in the presence of Pax2a in *hand2*-expressing cells (arrows). Stronger *hand2:egfp* expression in *hand2* mutants is consistent with our prior results (Perens et al., 2016). Scale bars: 100 μm (A-L,N-P), 50 μm (K’,L’,N’-P’).

To test whether sustained *osr1* expression would alter LVP development, we used the transgene *Tg*(*hsp70:osr1-t2A-BFP*) to overexpress *osr1* throughout the embryo. Strikingly, we found that induction of *osr1* overexpression at tailbud stage could inhibit LVP formation (Fig. 4G-J, Fig. S5). Additionally, *osr1* overexpression increased the formation of medial and proximal vessel progenitors, suggesting that *osr1* has distinct influences on the development of different subsets of vessel progenitor cells. Interestingly, *osr1* overexpression resulted in a range of severity for each of these vessel progenitor phenotypes, consistent with a dependence of vessel progenitor formation on precise *osr1* expression levels (Fig. S5).

We next examined whether there was a concomitant change in the IM when LVP formation was suppressed by *osr1* overexpression. Quantification of Pax2a^+^ cells showed a moderate, but significant, increase in the IM when *osr1* is overexpressed (Fig. 4K-M). What might be the origin of these additional IM cells? In *hand2* mutants, which exhibit IM expansion and loss of LVPs, a substantially increased number of Pax2a^+^ cells emerge within the *hand2*-expressing territory (Fig. 4P) (Perens et al., 2016). In contrast, in *osr1*-overexpressing embryos, Pax2a expression remained excluded from the *hand2*-expressing territory (Fig. 4O). Consistent with this difference, the increase in IM generated by *osr1* overexpression was less than that generated by *hand2* loss-of-function (Figs. 2I, 4M) (Perens et al., 2016). Together, our findings suggest that elevated *osr1* expression drives cells at the lateral IM border toward an IM fate, rather than a LVP fate. Like *hand2* loss-of-function, *osr1* overexpression can suppress LVP formation and increase IM production; but, unlike *hand2* loss-of-function, elevated *osr1* expression does not also convert the *hand2*-expressing lateral posterior mesoderm into Pax2a^+^ IM.

### *osr1* acts in opposition to *hand2* to promote intermediate mesoderm differentiation while inhibiting lateral vessel progenitor emergence

Altogether, our studies provide new insights into the roles of *osr1* in IM and vessel progenitor development. We show that *osr1* is both necessary and sufficient to promote the initial differentiation of some, but not all IM. Also, we reveal context-dependent roles for *osr1* in inhibiting vessel progenitor development, including an intriguing role in preventing the premature appearance of vessel progenitors at the lateral border of the IM. Finally, our findings suggest that the dynamic nature of *osr1* expression in the posterior mesoderm is necessary for balancing the extent of IM formation with the timing of neighboring LVP emergence.

How might *osr1* regulate both the IM and LVP lineages within the posterior mesoderm? Our findings indicate the presence of a unique territory at the boundary between the developing IM and the lateral mesoderm in which *osr1* initially acts in opposition to *hand2* to promote IM differentiation while inhibiting vessel progenitor differentiation; later, as *osr1* expression recedes, IM differentiation ceases and vessel progenitors emerge. Conceptually, we envision that the dynamic levels of *osr1* expression couple developmental timing with cell fate acquisition in order to set boundaries that delineate the extent of each progenitor territory. In future studies, it will be important to determine how directly or indirectly Osr1 and Hand2 regulate expression of the downstream genes that control IM and vessel identity, such as *pax2a* and *etv2*. Prior work suggested that Osr1 and its Drosophila homolog Odd function as transcriptional repressors (Goldstein et al., 2005; Tena et al., 2007). Considering the importance of reciprocal repressor interactions in establishing boundaries between neighboring progenitor territories in other developmental contexts (Briscoe and Small, 2015), it is interesting to speculate that, in the posterior mesoderm, a key function of Osr1 is to repress LVP formation while a primary role of Hand2 is to inhibit IM differentiation.

Our studies also suggest three subregions capable of contributing to the IM, arranged along the medial-lateral axis of zebrafish posterior mesoderm, with distinct genetic networks regulating IM formation in each area: a medial *osr1*-independent territory, a far lateral territory with latent IM-forming potential that is repressed by the sustained expression of *hand2*, and a boundary territory in between these in which a balance between *hand2* and *osr1* functions cell-autonomously to determine the precise amount and timing of IM and LVP formation. High-resolution lineage tracing will be necessary to delineate the precise fates of the cells in these different territories and to determine how their developmental potential is shaped by Osr1 and Hand2.

Over the long term, our understanding of the impact of *osr1* on medial-lateral patterning of the posterior mesoderm may have important implications for understanding congenital anomalies of the kidney and urinary tract (CAKUT), especially as *OSR1* mutations have been associated with a variety of CAKUT phenotypes, including renal hypoplasia and vesicoureteral reflux (Fillion et al., 2017; Zhang et al., 2011). Additionally, because generation of IM is a key step in the production of vascularized kidney organoids (Takasato et al., 2015), our in vivo analyses of *osr1* function during IM and vessel progenitor formation have the potential to inform future refinements of relevant in vitro differentiation protocols.

## MATERIALS AND METHODS

### Zebrafish

We generated embryos by breeding wild-type zebrafish, zebrafish heterozygous for the *osr1* mutant allele *osr1^el593^* (RRID: ZFIN_ZDB-ALT-171010-14) (Askary et al., 2017), zebrafish heterozygous for the *hand2* mutant allele *han^s6^* (RRID: ZFIN_ZDB-GENO-071003-2) (Yelon et al., 2000), and zebrafish carrying *Tg*(*hand2:egfp*)^*pd24*^ (RRID: ZFIN_ZDB-GENO-110128-35) (Kikuchi et al., 2011), *Tg*(*hsp70:osr1-t2A-BFP*)^*sd63*^, or *Tg*(*etv2:egfp*)^*ci1*^ (RRID: ZFIN_ZDB-GENO-110131-58) (Proulx et al., 2010). For induction of heat shock-regulated expression, embryos at tailbud stage were placed at 37°C for 1 hr and then returned to 28°C. Following heat shock, transgenic embryos were identified based on BFP fluorescence; embryos used for genotyping and cell counting in Figure 4 were confirmed to carry the transgene by PCR genotyping for the *bfp* coding region. Nontransgenic embryos were analyzed as controls. PCR genotyping was conducted as previously described for *osr1^el593^* mutants (Askary et al., 2017), for *han^s6^* mutants (Yelon et al., 2000), and for *han^s6^* mutants containing *Tg(hand2:EGFP)^pd24^* (Perens et al., 2016). All zebrafish work followed protocols approved by the UCSD IACUC.

### Creation of stable transgenic lines

To generate transgenes for heat-activated overexpression of *osr1,* we first amplified the *osr1* coding sequence from pCS2-*osr1* (Mudumana et al., 2008), using the primers 5’–AAAAAAGCAGGCTGCCACCGATGGGTAGTAAGACGCTC–3’ and 5’–CTCCTCCGGACCCGCCGCCGTACTTTATCTTGGCTGGC–3’, and cloned the amplicon into the vector *hsp70*-BamHI-t2a-BFP at the BamHI restriction site. We employed standard protocols to create transgenic founders (Fisher et al., 2006). The F1 progeny of prospective founder fish were screened for BFP fluorescence following heat shock for 1 hr at 37°C, and phenotypic analysis was performed on the F1 and F2 progeny of three separate founders carrying distinct integrations of *Tg*(*hsp70:osr1-t2A-BFP*). Similar phenotypes were observed in all three transgenic lines; data shown in Figures 4 and S5 depict results from the line *Tg*(*hsp70:osr1-t2A-BFP*)^*sd63*^.

### In situ hybridization

Standard whole-mount in situ hybridization were performed as previously described (Thomas et al., 2008) using the following probes: *atp1a1a.4* (ZDB-GENE-001212-4), *cdh17* (ZDB-GENE-030910-3), *etv2* (*etsrp*; ZDB-GENE-050622-14), *foxa2* (ZDB-GENE-980526-404), *gata1* (ZDB-GENE-980536-268), *hand2* (ZDB-GENE-000511-1), *lhx1a* (*lim1*; ZDB-GENE-980526-347), *osr1* (ZDB-GENE-070321-1), *pax2a* (ZDB-GENE-990415-8), *slc12a3* (ZDB-GENE-030131-9505), *slc20a1a* (ZDB-GENE-040426-2217), *sox17* (ZDB-GENE-991213-1), and *wt1b* (ZDB-GENE-050420-319).

### Immunofluorescence

Whole-mount immunofluorescence was performed as previously described (Cooke et al., 2005), using polyclonal antibodies against Pax2a at 1:100 dilution (Genetex, Irvine, CA; GTX128127) (RRID: AB_2630322), and against GFP at 1:250 dilution (Life Technologies, Carlsbad, CA; A10262) (RRID: AB_2534023), together with the secondary antibodies goat anti-chick Alexa Fluor 488 (Life Technologies, A11039) (RRID: AB_2534096) and goat anti-rabbit Alexa Fluor 594 (Life Technologies, A11012) (RRID: AB_10562717), both at 1:100 dilution. Samples were then placed in SlowFade Gold anti-fade reagent (Life Technologies) and mounted in 50% glycerol.

### Imaging

Bright-field images were captured with a Zeiss Axiocam on a Zeiss Axiozoom microscope and processed using Zeiss AxioVision. Confocal images were collected by a Leica SP5 or SP8 confocal laser-scanning microscope using a 10X dry objective and a slice thickness of 1 μm and analyzed using Imaris software (Bitplane, Switzerland).

### Cell counting

To count Pax2a^+^ cells, we flat-mounted and imaged embryos after dissecting away the yolk and the anterior portion of the embryo. We examined a representative 250 μm long region in roughly the middle of the IM, selecting contiguous regions that were unaffected by dissection artifacts. When the quality of the dissection allowed both the right and left sides of the embryo to be counted, we counted Pax2a^+^ cells on both sides and represented the embryo by the average of the two values; otherwise, only one side was counted. In all cases, Pax2a^+^ cells were identified through examination of both three-dimensional reconstructions and individual optical sections. To differentiate Pax2a^+^ cells from background immunofluorescence, we used Imaris to decrease brightness until staining clearly outside of the IM was no longer visible.

### Statistics and replication

Statistical analysis of data was performed using Graphpad Prism 8 to conduct non-parametric Mann-Whitney U tests. All results represent at least two independent experiments (technical replicates) in which multiple embryos, from multiple independent matings, were analyzed (biological replicates). For wild-type and mutant in situ hybridization results, phenotypes shown are representative examples from at least 10 embryos for wild-type and *osr1* mutant phenotypes and at least 5 embryos for *hand2* mutant and *hand2*; *osr1* double mutant phenotypes. For wild-type, transgenic and mutant antibody staining results for which the phenotype was not quantified (Fig. 4I,J,N-P), phenotypes shown are representative examples from at least 5 embryos.

## ACKNOWLEDGEMENTS

We thank members of the Yelon lab for valuable discussions, A. Houk for providing the *hsp70*-*BamHI-t2A-BFP* plasmid, H. Kwan for generating the *hsp70*:*osr1-t2A-BFP* plasmid, and T. Sanchez and the UCSD Animal Care Program for zebrafish care.

## AUTHOR CONTRIBUTIONS

Conceptualization: E.A.P, D.Y.; Methodology: E.A.P, J.T.D., D.Y.; Formal analysis: E.A.P, J.T.D., D.Y.; Investigation: E.A.P, J.T.D., A.Q.; Resources: A.A., J.G.C.; Writing – original draft: E.A.P, D.Y.; Writing – review & editing: E.A.P, J.T.D., A.Q., A.A., J.G.C, D.Y.; Supervision: J.G.C., D.Y.; Project administration: D.Y.; Funding acquisition: E.A.P, J.G.C., D.Y.

## SUPPLEMENTARY FIGURE LEGENDS

**Supplementary Figure S1.**
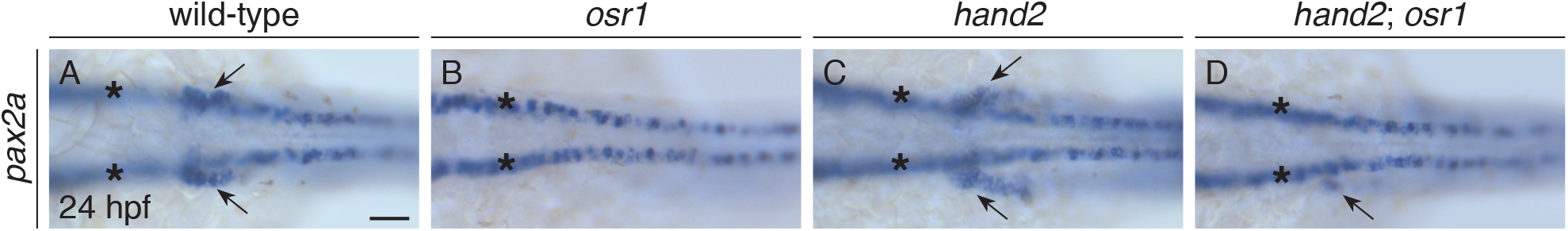
Proximal pronephron defects in *osr1* mutants are partially suppressed by *hand2* loss-of-function. (A-D) Dorsal views, anterior to the left, of wild-type (A), *osr1* mutant (B), *hand2* mutant (C), and *hand2*; *osr1* double mutant (D) embryos at 26 hpf. In situ hybridization shows *pax2a* expression in the glomerular precursors and the neck region (arrows) of the pronephron. Compared to wild-type (A), *pax2a* expression is absent in *osr1* mutants (B), expanded in *hand2* mutants (C), and most similar to wild-type in *hand2*; *osr1* double mutants (D). Expression in overlying spinal neurons (asterisks) is unaffected. Scale bar: 25 μm.

**Supplementary Figure S2.**
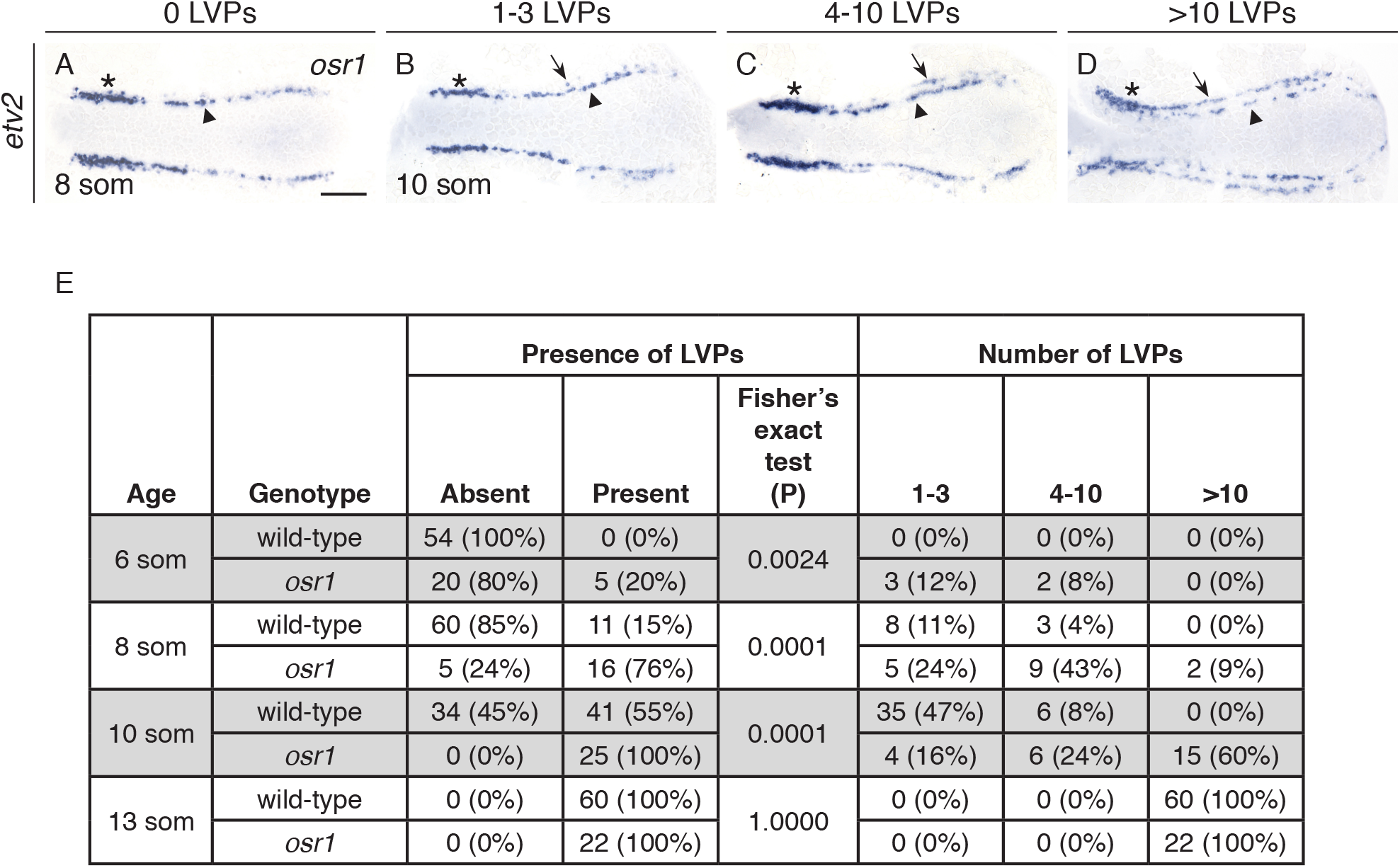
Premature lateral vessel progenitors in *osr1* mutant embryos. (A-D) Dorsal views, anterior to the left, of *osr1* mutant embryos at 8 som (A) and 10 som (B-D). In situ hybridization demonstrates that *etv2* is expressed bilaterally in medial (arrowheads) and lateral (arrows) territories. Representative examples are shown for embryos with 0 LVPs (A), 1-3 LVPs (B), 4-10 LVPs (C), and >10 LVPs (D). Embryos were categorized based on the number of cells observed on whichever side of the mesoderm exhibited more LVPs. Note the increased *etv2* expression in the most proximal portion of the posterior mesoderm (asterisks); this phenotype was also observed previously in *osr1* morphants (Mudumana et al., 2008). (E) Number of embryos and corresponding percentages in each category, as summarized in Fig. 3E. At each stage, the proportions of embryos with and without LVPs were compared between *osr1* mutants and their wild-type siblings using Fisher’s exact test; P values are provided for each comparison. At 8 som and 10 som, and to a lesser degree at 6 som, we observed significantly more *osr1* mutant embryos with LVPs than wild-type siblings with LVPs. Scale bar: 100 μm.

**Supplementary Figure S3.**
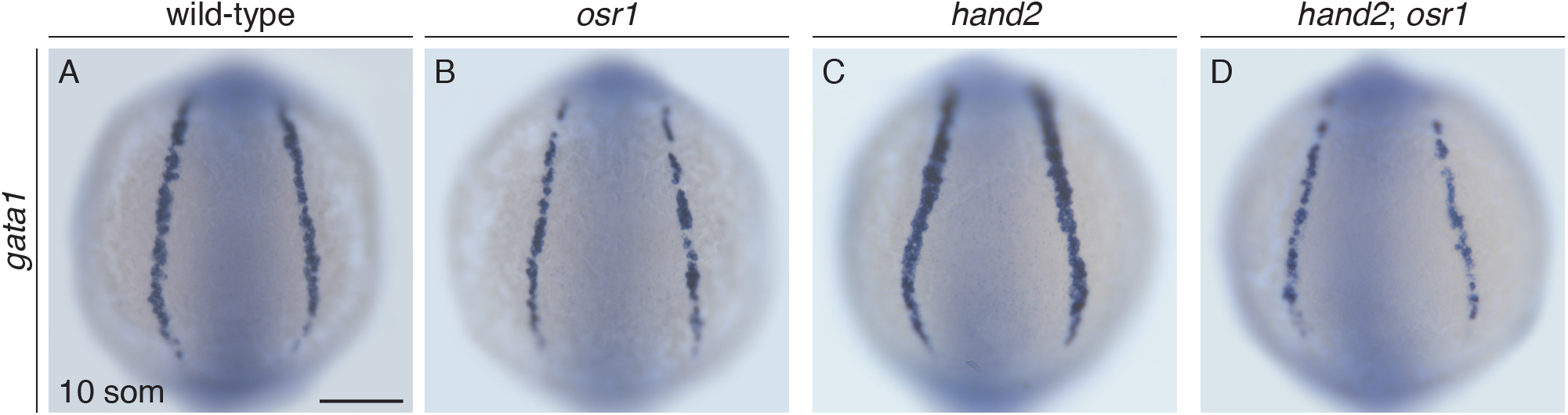
*hand2* loss-of-function does not suppress the blood precursor defects in *osr1* mutants. (A-D) Dorsal views, anterior to the top, of wild-type (A), *osr1* mutant (B), *hand2* mutant (C), and *hand2*; *osr1* double mutant embryos (D) at 10 som. In situ hybridization shows *gata1* expression in blood precursors. Compared to wild-type (A), *gata1* expression is decreased throughout the posterior mesoderm in *osr1* mutants (B), unchanged in *hand2* mutants (C), and relatively similar to *osr1* mutants in *hand2*; *osr1* double mutant embryos (D). In contrast, *osr1* morphants were shown to exhibit reduction in *gata1* expression only at the proximal end of the *gata1* expression domain (Mudumana et al., 2008). Scale bar: 100 μm.

**Supplementary Figure S4.**
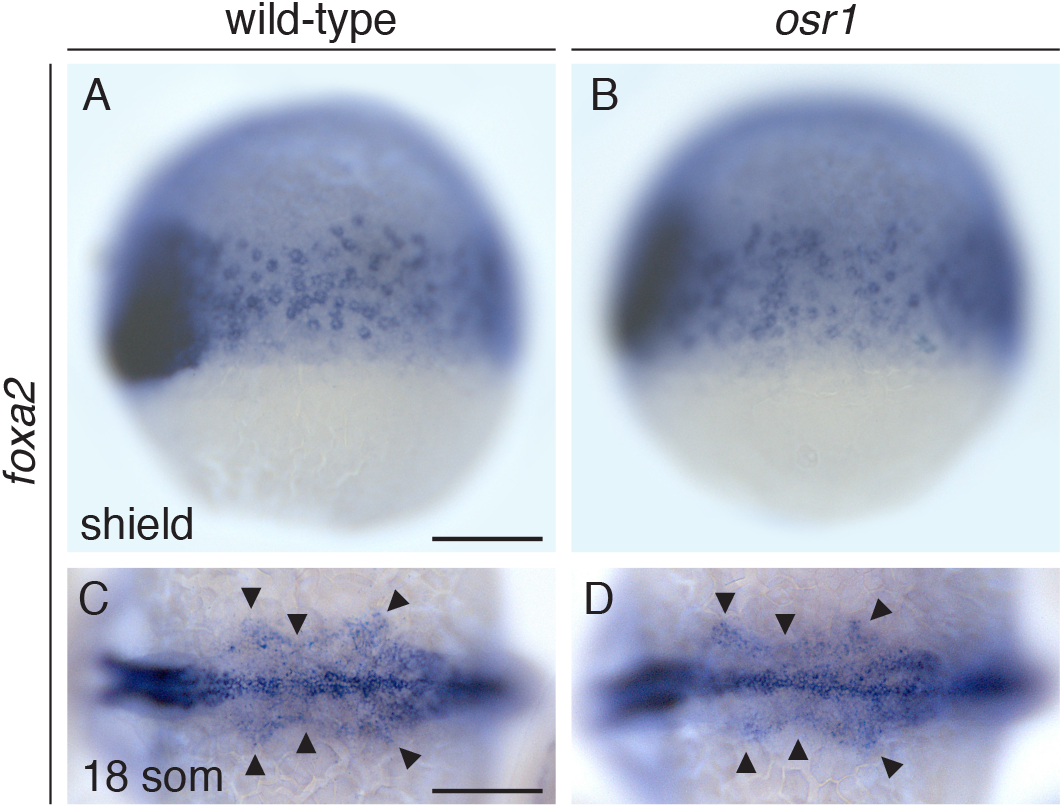
Endoderm formation is grossly normal in *osr1* mutant embryos. (A-D) Lateral views, dorsal to the left, at shield stage (A,B) and dorsal views, anterior to the left, at 18 som (C,D) in wild-type (A,C) and *osr1* mutant (B,D) embryos. In situ hybridization shows *foxa2* expression in the embryonic shield and endodermal precursors (A,B) and in the pharyngeal endoderm (arrowheads) (C,D). We did not find any consistent differences in the endoderm when comparing wild-type and *osr1* mutant embryos at either shield stage or 18 som. At shield stage, we did observe a varied amount of *foxa2* and *sox17* (data not shown) expression among the embryos, but the overall distribution of expression was comparable in wild-type and *osr1* mutants. In contrast, *osr1* morphants were reported to exhibit increased numbers of endodermal cells expressing *foxa2* and *sox17* (Mudumana et al., 2008; Terashima et al., 2014); we cannot rule out contributions of maternal effect or genetic compensation to the phenotype of *osr1* mutants. Scale bars: 100 μm.

**Supplementary Figure S5.**
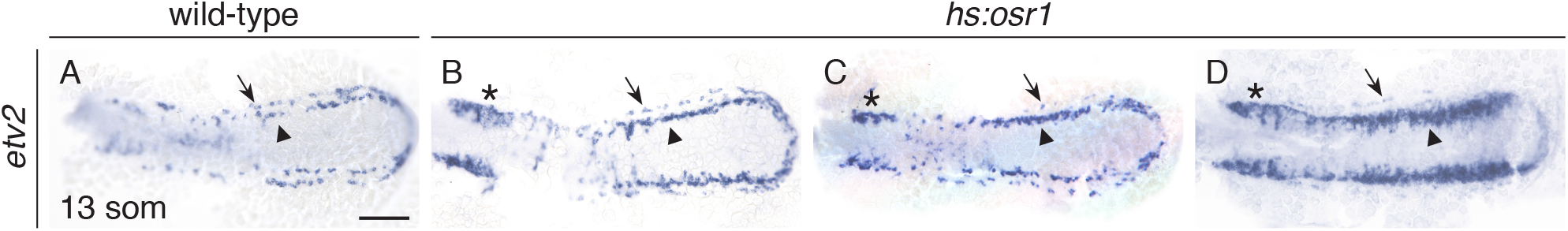
Overexpression of *osr1* yields a range of vessel progenitor phenotypes. (A-D) Dorsal views, anterior to the left, of wild-type (A) and *hs:osr1* embryos (B-D) at 13 som. (A) In situ hybridization demonstrates that *etv2* is expressed in medial (arrowheads), lateral (arrows), and proximal (asterisk) territories within the wild-type posterior mesoderm. (B-D) In *hs:osr1* embryos, a range of phenotypes was observed, including embryos with mildly increased medial expression and mildly reduced lateral expression (B), embryos with moderately increased medial expression and greatly reduced lateral expression (C), and embryos with a greatly expanded territory of medial expression and absent lateral expression (D). We could not distinguish whether the suppression of LVP formation was due to a delay of LVP formation or a complete inability to form LVPs; after 14 som, the vessel progenitors migrate toward the midline and the medial and lateral vessel progenitors become indistinguishable, as there are no known molecular markers that distinguish the two populations (Kohli et al., 2013). Surprisingly, the proximal *etv2* expression territory appeared expanded in *hs:osr1* embryos (B-D, asterisks), similar to the *osr1* mutant phenotype (Fig. 3D,G). Taken together, the divergent responses of each of the *etv2*-expressing territories to *osr1* loss-of-function and gain-of-function suggest that *osr1* plays a different, and possibly independent, role in the regulation of each vessel progenitor territory. It is intriguing to consider whether these differences in regulation correlate with the subpopulations of *etv2*-expressing cells distinguished by unique transcriptional signatures, as determined by single-cell RNA-sequencing (Chestnut et al., 2020). Scale bar: 100 μm.

## REFERENCES

Askary, A., Xu, P., Barske, L., Bay, M., Bump, P., Balczerski, B., Bonaguidi, M.A., Crump, J.G., 2017. Genome-wide analysis of facial skeletal regionalization in zebrafish. Development 144, 2994–3005.

Barnes, J.D., Crosby, J.L., Jones, C.M., Wright, C.V., Hogan, B.L., 1994. Embryonic expression of Lim-1, the mouse homolog of Xenopus Xlim-1, suggests a role in lateral mesoderm differentiation and neurogenesis. Dev Biol 161, 168–178.

Briscoe, J., Small, S., 2015. Morphogen rules: design principles of gradient-mediated embryo patterning. Development 142, 3996–4009.

Carroll, T.J., Vize, P.D., 1999. Synergism between Pax-8 and lim-1 in embryonic kidney development. Dev Biol 214, 46–59.

Carroll, T.J., Wallingford, J.B., Vize, P.D., 1999. Dynamic patterns of gene expression in the developing pronephros of Xenopus laevis. Dev Genet 24, 199–207.

Chestnut, B., Casie Chetty, S., Koenig, A.L., Sumanas, S., 2020. Single-cell transcriptomic analysis identifies the conversion of zebrafish Etv2-deficient vascular progenitors into skeletal muscle. Nat Commun 11, 2796.

Cirio, M.C., Hui, Z., Haldin, C.E., Cosentino, C.C., Stuckenholz, C., Chen, X., Hong, S.K., Dawid, I.B., Hukriede, N.A., 2011. Lhx1 is required for specification of the renal progenitor cell field. PLoS One 6, e18858.

Cooke, J.E., Kemp, H.A., Moens, C.B., 2005. EphA4 is required for cell adhesion and rhombomere-boundary formation in the zebrafish. Curr Biol 15, 536–542.

Davidson, A.J., Lewis, P., Przepiorski, A., Sander, V., 2019. Turning mesoderm into kidney. Semin Cell Dev Biol 91, 86–93.

Dressler, G.R., 2009. Advances in early kidney specification, development and patterning. Development 136, 3863–3874.

Dressler, G.R., Deutsch, U., Chowdhury, K., Nornes, H.O., Gruss, P., 1990. Pax2, a new murine paired-box-containing gene and its expression in the developing excretory system. Development 109, 787–795.

Fillion, M.L., El Andalousi, J., Tokhmafshan, F., Murugapoopathy, V., Watt, C.L., Murawski, I.J., Capolicchio, J.P., El-Sherbiny, M., Jednak, R., Gupta, I.R., 2017. Heterozygous loss-of-function mutation in Odd-skipped related 1 (Osr1) is associated with vesicoureteric reflux, duplex systems, and hydronephrosis. Am J Physiol Renal Physiol 313, F1106–F1115.

Fisher, S., Grice, E.A., Vinton, R.M., Bessling, S.L., Urasaki, A., Kawakami, K., McCallion, A.S., 2006. Evaluating the biological relevance of putative enhancers using Tol2 transposon-mediated transgenesis in zebrafish. Nat Protoc 1, 1297–1305.

Gerlach, G.F., Wingert, R.A., 2013. Kidney organogenesis in the zebrafish: insights into vertebrate nephrogenesis and regeneration. Wiley Interdiscip Rev Dev Biol 2, 559–585.

Goldstein, R.E., Cook, O., Dinur, T., Pisante, A., Karandikar, U.C., Bidwai, A., Paroush, Z., 2005. An eh1-like motif in odd-skipped mediates recruitment of Groucho and repression in vivo. Mol Cell Biol 25, 10711–10720.

James, R.G., Kamei, C.N., Wang, Q., Jiang, R., Schultheiss, T.M., 2006. Odd-skipped related 1 is required for development of the metanephric kidney and regulates formation and differentiation of kidney precursor cells. Development 133, 2995–3004.

Kikuchi, K., Holdway, J.E., Major, R.J., Blum, N., Dahn, R.D., Begemann, G., Poss, K.D., 2011. Retinoic acid production by endocardium and epicardium is an injury response essential for zebrafish heart regeneration. Dev Cell 20, 397–404.

Kohli, V., Schumacher, J.A., Desai, S.P., Rehn, K., Sumanas, S., 2013. Arterial and venous progenitors of the major axial vessels originate at distinct locations. Dev Cell 25, 196–206.

Kroeger, P.T., Jr., Drummond, B.E., Miceli, R., McKernan, M., Gerlach, G.F., Marra, A.N., Fox, A., McCampbell, K.K., Leshchiner, I., Rodriguez-Mari, A., BreMiller, R., Thummel, R., Davidson, A.J., Postlethwait, J., Goessling, W., Wingert, R.A., 2017. The zebrafish kidney mutant zeppelin reveals that brca2/fancd1 is essential for pronephros development. Dev Biol 428, 148–163.

Lun, K., Brand, M., 1998. A series of no isthmus (noi) alleles of the zebrafish pax2.1 gene reveals multiple signaling events in development of the midbrain-hindbrain boundary. Development 125, 3049–3062.

Mudumana, S.P., Hentschel, D., Liu, Y., Vasilyev, A., Drummond, I.A., 2008. odd skipped related1 reveals a novel role for endoderm in regulating kidney versus vascular cell fate. Development 135, 3355–3367.

Mugford, J.W., Sipila, P., McMahon, J.A., McMahon, A.P., 2008. Osr1 expression demarcates a multi-potent population of intermediate mesoderm that undergoes progressive restriction to an Osr1-dependent nephron progenitor compartment within the mammalian kidney. Dev Biol 324, 88–98.

Perens, E.A., Garavito-Aguilar, Z.V., Guio-Vega, G.P., Pena, K.T., Schindler, Y.L., Yelon, D., 2016. Hand2 inhibits kidney specification while promoting vein formation within the posterior mesoderm. Elife 5. e19941.

Proulx, K., Lu, A., Sumanas, S., 2010. Cranial vasculature in zebrafish forms by angioblast cluster-derived angiogenesis. Dev Biol 348, 34–46.

Prummel, K.D., Nieuwenhuize, S., Mosimann, C., 2020. The lateral plate mesoderm. Development 147: dev175059.

Schulte-Merker, S., Stainier, D.Y., 2014. Out with the old, in with the new: reassessing morpholino knockdowns in light of genome editing technology. Development 141, 3103–3104.

Takasato, M., Er, P.X., Chiu, H.S., Maier, B., Baillie, G.J., Ferguson, C., Parton, R.G., Wolvetang, E.J., Roost, M.S., Chuva de Sousa Lopes, S.M., Little, M.H., 2015. Kidney organoids from human iPS cells contain multiple lineages and model human nephrogenesis. Nature 526, 564–568.

Tena, J.J., Neto, A., de la Calle-Mustienes, E., Bras-Pereira, C., Casares, F., Gomez-Skarmeta, J.L., 2007. Odd-skipped genes encode repressors that control kidney development. Dev Biol 301, 518–531.

Terashima, A.V., Mudumana, S.P., Drummond, I.A., 2014. Odd skipped related 1 is a negative feedback regulator of nodal-induced endoderm development. Dev Dyn 243, 1571–1580.

Thomas, N.A., Koudijs, M., van Eeden, F.J., Joyner, A.L., Yelon, D., 2008. Hedgehog signaling plays a cell-autonomous role in maximizing cardiac developmental potential. Development 135, 3789–3799.

Tomar, R., Mudumana, S.P., Pathak, N., Hukriede, N.A., Drummond, I.A., 2014. osr1 Is Required for Podocyte Development Downstream of wt1a. J Am Soc Nephrol 11, 2539–2545.

Torres, M., Gomez-Pardo, E., Dressler, G.R., Gruss, P., 1995. Pax-2 controls multiple steps of urogenital development. Development 121, 4057–4065.

Tsang, T.E., Shawlot, W., Kinder, S.J., Kobayashi, A., Kwan, K.M., Schughart, K., Kania, A., Jessell, T.M., Behringer, R.R., Tam, P.P., 2000. Lim1 activity is required for intermediate mesoderm differentiation in the mouse embryo. Dev Biol 223, 77–90.

Wang, Q., Lan, Y., Cho, E.S., Maltby, K.M., Jiang, R., 2005. Odd-skipped related 1 (Odd 1) is an essential regulator of heart and urogenital development. Dev Biol 288, 582–594.

Yelon, D., Ticho, B., Halpern, M.E., Ruvinsky, I., Ho, R.K., Silver, L.M., Stainier, D.Y., 2000. The bHLH transcription factor hand2 plays parallel roles in zebrafish heart and pectoral fin development. Development 127, 2573–2582.

Zhang, Z., Iglesias, D., Eliopoulos, N., El Kares, R., Chu, L., Romagnani, P., Goodyer, P., 2011. A variant OSR1 allele which disturbs OSR1 mRNA expression in renal progenitor cells is associated with reduction of newborn kidney size and function. Hum Mol Genet 20, 4167–4174.

